# Identification of potential biomarkers or therapeutic targets of mesenchymal stem cells in multiple myeloma by bioinformatics analysis

**DOI:** 10.1101/2020.06.16.153676

**Authors:** Zhi-Ran Li, Wen-Ke Cai, Qin Yang, Ming-Li Shen, Hua-Zhu Zhang, Qian Huang, Gui-Xin Zhao, Ke-Yan Chen, Gong-Hao He

## Abstract

**Objectives:** Mesenchymal stem cells (MSCs) play important roles in multiple myeloma (MM) pathogenesis. Previous studies have discovered a group of MM-associated potential biomarkers in MSCs derived from bone marrow (BM-MSCs). However, no study of the bioinformatics analysis was conducted to explore the key genes and pathways of MSCs derived from adipose (AD-MSCs) in MM. The aim of this study was to screen potential biomarkers or therapeutic targets of AD-MSCs and BM-MSCs in MM.

**Methods:** The gene expression profiles of AD-MSCs (GSE133346) and BM-MSCs (GSE36474) were downloaded from Gene Expression Omnibus (GEO) database. Gene Oncology (GO) enrichment, Kyoto Encyclopedia of Genes and Genomes (KEGG) pathway analysis and protein-protein interaction (PPI) network of differentially expressed genes (DEGs) were performed.

**Results:** A total of 456 common downregulated DEGs in two datasets were identified and the remaining DEGs in GSE133346 were further identified as specific DEGs of AD-MSCs. Furthermore, a PPI network of common downregulated DEGs was constructed and seven hub genes were identified. Importantly, cell cycle was the most significantly enrichment pathway both in AD-MSCs and BM-MSCs from MM patients.

**Conclusion:** We identified key genes and pathways closely related with MM progression, which may act as potential biomarkers or therapeutic targets of MM.

## 1 Introduction

Multiple myeloma (MM) is a severe cancer of plasma cells and is characterized by unlimited proliferation of malignant plasma cells in the bone marrow [1]. It represents the second most common hematologic malignancy, inflicting approximately 13,000 mortalities in the United States in 2018 [2]. Unfortunately, the exact molecular mechanism underlying the development of MM remains unclear. Recently, it was indicated that mesenchymal stem cells (MSCs) play important roles in MM pathogenesis, which have both tumor supportive and inhibitory effects in MM [3-5]. However, further research is still needed to investigate the mechanism of MSCs in MM development and progression [3], which holds promise for finding potential drug targets and diagnostic biomarkers of MM.

MSCs reside within the connective tissue of virtually all organs, including bone marrow, muscle, thymus and adipose tissue as well [6, 7]. Nevertheless, previous studies mainly focused on bone marrow-derived mesenchymal stem cells (BM-MSCs) and have discovered a group of MM-associated potential biomarkers and key pathways by gene expression profiling analysis with microarray technology [8, 9]. However, no study of the bioinformatics analysis was conducted in other MSCs due to lacking of relevant microarray datasets. Recently, a new microarray dataset (GSE133346) [10] of adipose-derived mesenchymal stem cells (AD-MSCs) was uploaded to the Gene Expression Omnibus (GEO) database, which offers reliable data to a systematic bioinformatics analysis of AD-MSCs and is worthy of further research to better understand the role of AD-MSCs in the pathogenesis of MM. In addition, comparing and analyzing the DEGs between AD-MSCs and BM-MSCs from the same platform would also provide more information for identifying potential specific markers of AD-MSC in MM.

Based on this background, GSE133346, along with the dataset (GSE36474) [11] of BM-MSCs deriving from the same platform, were downloaded from the GEO database for further analysis in the current study. Based on these 2 datasets, a joint bioinformatics analysis of two gene expression profiles was performed to identify MM-associated DEGs. Besides, Gene Ontology (GO) term enrichment analysis, Kyoto Encyclopedia of Genes and Genomes (KEGG) pathway analysis, and protein-protein interaction (PPI) network analysis were applied to annotate gene function and screen hub genes. The objective of this study was to identify potential crucial genes and key pathways closely related to MM both in AD-MSCs and BM-MSCs. The results of the present study may improve our understanding of the molecular mechanisms underlying MM and contribute to finding specific biomarkers of AD-MSCs in MM.

## 2 Materials and methods

### 2.1 Microarray data information and DEGs

Data of gene chips GSE133346, GSE36474 were downloaded from the GEO database (http://www.ncbi.nlm.nih.gov/geo/). Both datasets were based on the microarray platform of Affymetrix Human Genome U133 Plus 2.0 Array (GPL570 platform) to avoid the impact of different detection platforms on the analysis. The gene expression profile GSE133346 included AD-MSCs samples from 12 MM patients and 12 healthy controls. The GSE36474 included BM-MSCs samples from 4 MM patients and 3 healthy controls.

### 2.2 Identification of DEGs

The raw data in CEL files were preprocessed, including background correction, normalization and Log_2_ conversion via the affy package in R software (version 3.6.0). The limma package of R was used to screen DEGs from GEO datasets. Log_2_-fold change (log_2_FC) was reckoned to identify genes with expression-level differences. The genes meeting the cut-off criteria of adjusted P < 0.05 and |log_2_FC| > 1 were selected as DEGs.

### 2.3 Functional enrichment analysis

GO analysis was carried out with the R package ggplot2. GO categories are classified into 3 groups: biological process (BP), cellular component (CC), and molecular function (MF). Another online biological tool, KOBAS 3.0 (http://kobas.cbi.pku.edu.cn), was used for the KEGG pathway functional enrichment. P < 0.05 was considered to be statistically significant.

### 2.4 PPI network construction and analysis of modules

The STRING database (available online: http://string-db.org) was used to obtain the interactions between proteins. This study performed the PPI network using overlapped DEGs in the two independent datasets (GSE133346 and GSE36474). The cut-off criterion was set as Confidence score > 0.9. Then, the network was visualized in Cytoscape (version 3.7.1). The hub genes were identified by Cytohubba with three ranking methods, including Degree, Maximum Neighborhood Component (MNC), and Maximal Clique Centrality (MCC). Any overlaps in the top 10 genes from three ranking methods were defined as hub genes. The Molecular Complex Detection (MCODE) in Cytoscape software was used to explore the significant modules of the PPI network. Moreover, genes in selected modules were analyzed via KOBAS to examine KEGG pathway enrichment analyses. P < 0.05 was considered significantly enrichment.

## 3 Results

### 3.1 Identification of DEGs in MM

There were 5605 DEGs (6 upregulated and 5599 downregulated) in GSE133346 and 1132 DEGs (139 upregulated and 993 downregulated) in GSE36474, respectively (Fig. 1A and B). Intersection analysis of the two datasets identified 456 common downregulated DEGs (Fig. 1B). For GSE133346 dataset, after removing 456 overlapped DEGs, the remaining genes were further identified as specific DEGs of AD-MSCs (6 specifically upregulated and 5143 specifically downregulated) (Fig. 1A and B). For GSE36474 dataset, after removing 456 overlapped DEGs, the rest were further identified as specific DEGs of BM-MSCs (139 specifically upregulated and 537 specifically downregulated) (Fig. 1A and B). To visualize the differential expression levels in two microarray expression profiles following standardization, the volcano plots were used for DEGs in GSE133346 (Fig. 1C) and GSE36474 (Fig. 1D).

**Fig. 1.**
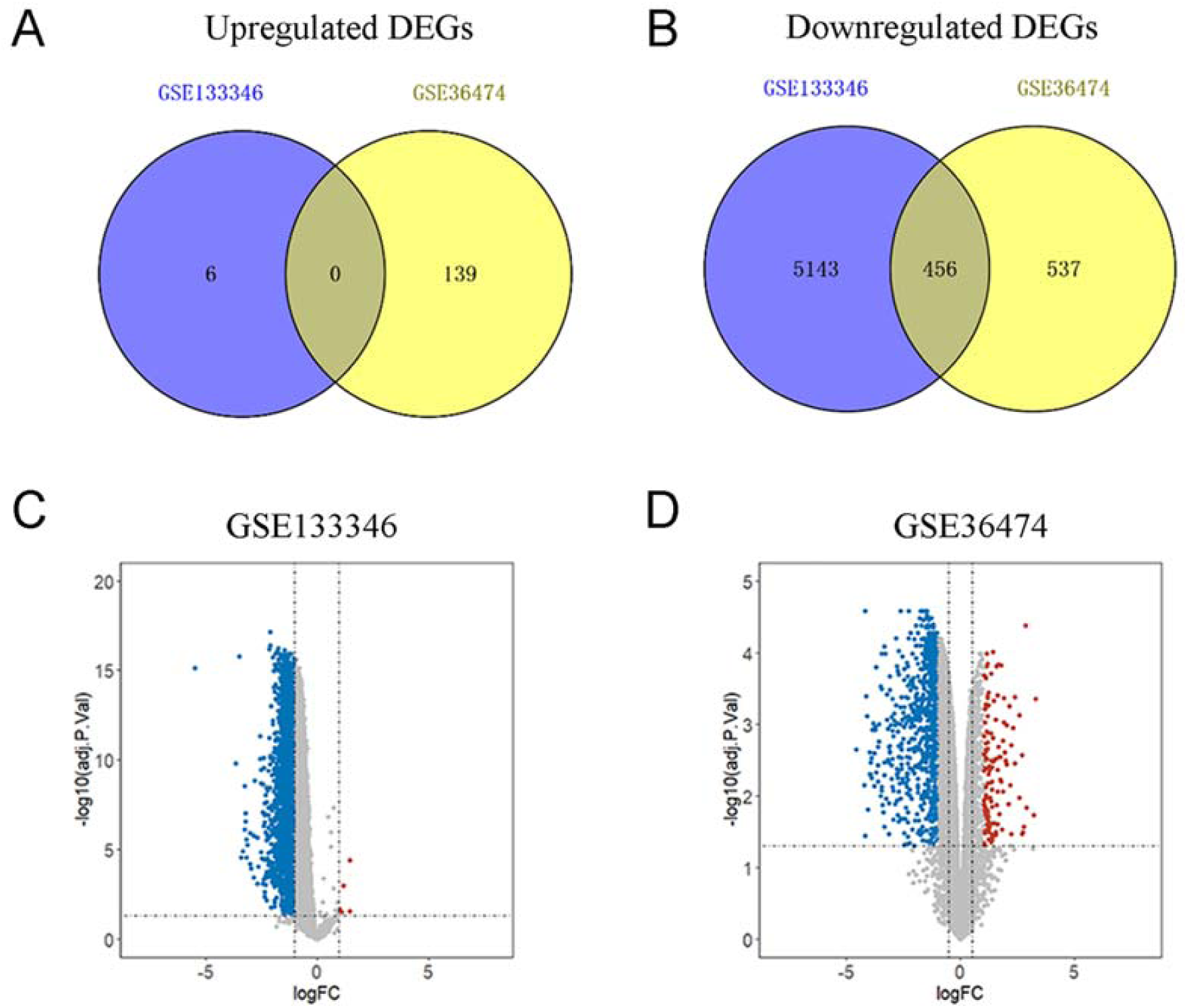
Identification of differentially expressed genes (DEGs) in two profile datasets. Venn analysis of DEGs: (A) Common upregulated DEGs, (B) common downregulated DEGs. Respective volcano plot of the two datasets: (C) GSE133346, (D) GSE36474. The red points represent upregulated genes with log_2_FC > 1 and adjusted P < 0.05. The blue points represent downregulated genes with log_2_FC < −1 and adjusted p < 0.05.

### 3.2 DEGs ontology analysis in MM

To determine the biological functions of DEGs, GO analysis was applied using the ggplot2 R package. For 456 common downregulated DEGs, the top 10 most significant terms of BP, CC, and MF are shown in Table 1. The GO analysis results showed that the DEGs were particularly enriched in BP, including DNA replication, nuclear division and organelle fission. With regard to CC, the common downregulated DEGs were significantly enriched in chromosomal region, spindle, and centrosome. In addition, the common downregulated DEGs in MF were mainly enriched in chromatin binding, catalytic activity, acting on DNA, and ATPase activity. These results showed that most of the overlapped DEGs were significantly enriched in GO terms related to cell proliferation.

**Table 1.** GO enrichment analysis of common downregulated DEGs.

| Category | Term | Description | Count | P value |
| --- | --- | --- | --- | --- |
| BP | GO:0006260 | DNA replication | 53 | 8.26E-39 |
| BP | GO:0140014 | Mitotic nuclear division | 46 | 1.04E-32 |
| BP | GO:0006261 | DNA-dependent DNA replication | 36 | 1.22E-31 |
| BP | GO:0000280 | Nuclear division | 51 | 1.70E-29 |
| BP | GO:0048285 | Organelle fission | 51 | 2.28E-27 |
| BP | GO:0000819 | Sister chromatid segregation | 35 | 3.08E-27 |
| BP | GO:0007059 | Chromosome segregation | 43 | 1.41E-26 |
| BP | GO:0000070 | Mitotic sister chromatid segregation | 32 | 3.23E-26 |
| BP | GO:0098813 | Nuclear chromosome segregation | 38 | 3.41E-25 |
| BP | GO:0044786 | Cell cycle DNA replication | 22 | 2.15E-23 |
| CC | GO:0098687 | Chromosomal region | 46 | 1.00E-25 |
| CC | GO:0005819 | spindle | 41 | 1.23E-21 |
| CC | GO:0000793 | Condensed chromosome | 33 | 2.16E-21 |
| CC | GO:0000775 | chromosome, centromeric region | 29 | 3.08E-18 |
| CC | GO:0000776 | kinetochore | 23 | 7.94E-16 |
| CC | GO:0000779 | Condensed chromosome, centromeric region | 21 | 2.52E-15 |
| CC | GO:0000777 | Condensed chromosome kinetochore | 20 | 4.57E-15 |
| CC | GO:0000922 | Spindle pole | 23 | 2.26E-14 |
| CC | GO:0005813 | centrosome | 39 | 2.58E-14 |
| CC | GO:0005657 | Replication fork | 13 | 2.46E-11 |
| MF | GO:0003697 | single-stranded DNA binding | 12 | 3.70E-07 |
| MF | GO:0003684 | Damaged DNA binding | 10 | 4.21E-07 |
| MF | GO:0140097 | Catalytic activity, acting on DNA | 15 | 7.16E-07 |
| MF | GO:0043142 | single-stranded DNA-dependent ATPase activity | 5 | 2.73E-06 |
| MF | GO:0035173 | Histone kinase activity | 5 | 6.68E-06 |
| MF | GO:0008094 | DNA-dependent ATPase activity | 9 | 1.85E-05 |
| MF | GO:0003682 | Chromatin binding | 22 | 1.41E-04 |
| MF | GO:0004386 | Helicase activity | 10 | 3.61E-04 |
| MF | GO:0042623 | ATPase activity, coupled | 13 | 7.04E-04 |
| MF | GO:0016887 | ATPase activity | 15 | 8.57E-04 |

GO: Gene Ontology; DEGs: differentially expressed genes; BP: biological process; CC: cell component; MF: molecular function.

For AD-MSCs specific DEGs in GSE133346, a total of 6 DEGs (OLR1, AMIGO2, GPR116, BEX1, CNN1, INHBA) were found to be upregulated with only one gene (INHBA) enriched in 2 MF GO terms (i.e., transmembrane receptor protein serine/threonine kinase binding and receptor serine/threonine kinase binding) (Table 2). The 5143 specifically downregulated DEGs in AD-MSCs in BP were mostly enriched in GO terms related to metabolic process, such as positive regulation of catabolic process, positive regulation of cellular catabolic process and fatty acid metabolic process. With regard to CC and MF, these downregulated DEGs were significantly enriched in endosome membrane, early endosome and nuclear envelope (for CC) and in ubiquitin-like protein transferase activity (for MF), respectively (Table 2). For BM-MSCs specific DEGs in GSE36474, the most significant terms are shown in Supplementary Table 1.

**Table 2.**
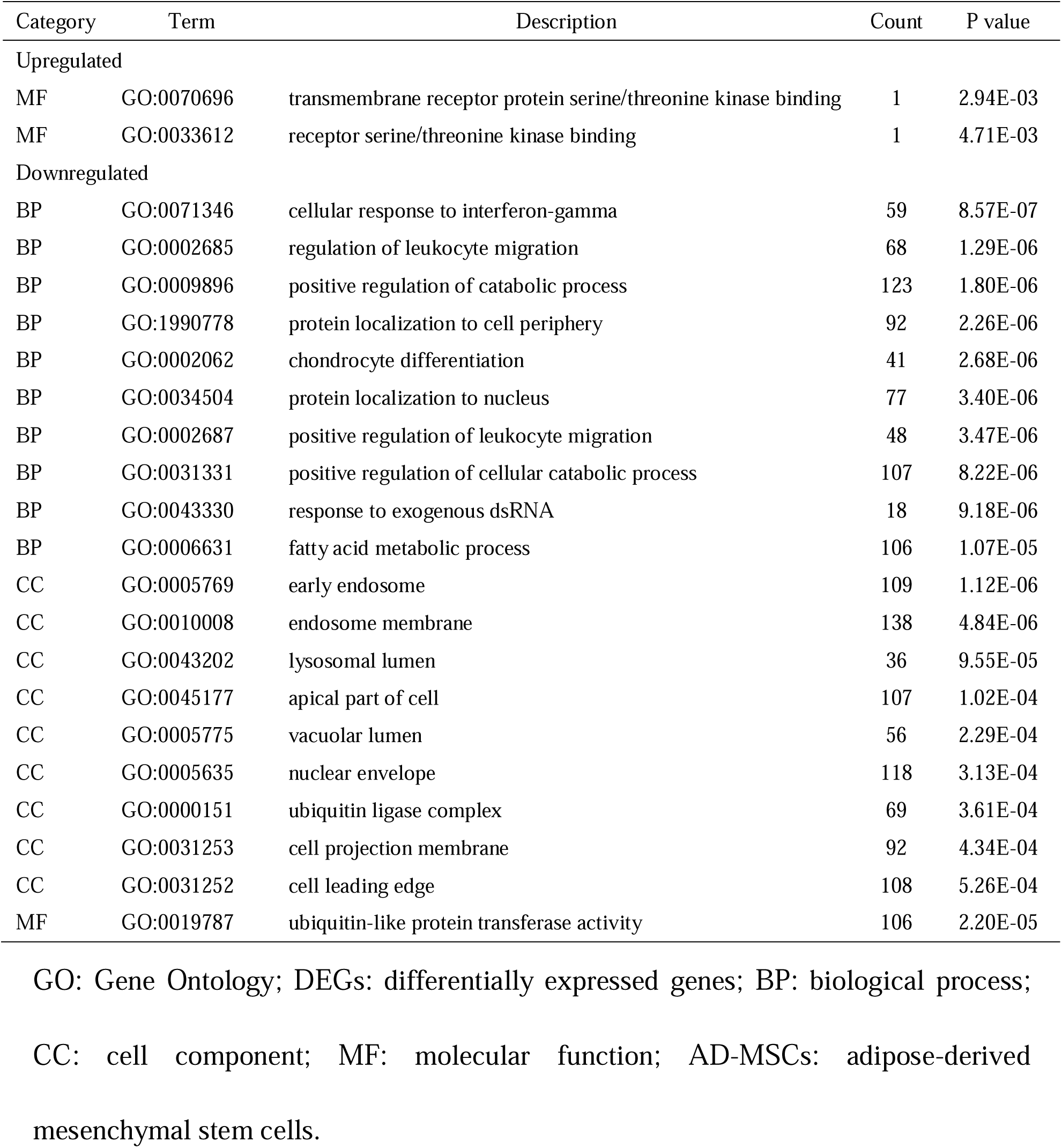
GO enrichment analysis of the specifically up- and downregulated DEGs in AD-MSCs.

| Category | Term | Description | Count | P value |
| --- | --- | --- | --- | --- |
| Upregulated |  |  |  |  |
| MF | GO:0070696 | transmembrane receptor protein serine/threonine kinase binding | 1 | 2.94E-03 |
| MF | GO:0033612 | receptor serine/threonine kinase binding | 1 | 4.71E-03 |
| Downregulated |  |  |  |  |
| BP | GO:0071346 | cellular response to interferon-gamma | 59 | 8.57E-07 |
| BP | GO:0002685 | regulation of leukocyte migration | 68 | 1.29E-06 |
| BP | GO:0009896 | positive regulation of catabolic process | 123 | 1.80E-06 |
| BP | GO:1990778 | protein localization to cell periphery | 92 | 2.26E-06 |
| BP | GO:0002062 | chondrocyte differentiation | 41 | 2.68E-06 |
| BP | GO:0034504 | protein localization to nucleus | 77 | 3.40E-06 |
| BP | GO:0002687 | positive regulation of leukocyte migration | 48 | 3.47E-06 |
| BP | GO:0031331 | positive regulation of cellular catabolic process | 107 | 8.22E-06 |
| BP | GO:0043330 | response to exogenous dsRNA | 18 | 9.18E-06 |
| BP | GO:0006631 | fatty acid metabolic process | 106 | 1.07E-05 |
| CC | GO:0005769 | early endosome | 109 | 1.12E-06 |
| CC | GO:0010008 | endosome membrane | 138 | 4.84E-06 |
| CC | GO:0043202 | lysosomal lumen | 36 | 9.55E-05 |
| CC | GO:0045177 | apical part of cell | 107 | 1.02E-04 |
| CC | GO:0005775 | vacuolar lumen | 56 | 2.29E-04 |
| CC | GO:0005635 | nuclear envelope | 118 | 3.13E-04 |
| CC | GO:0000151 | ubiquitin ligase complex | 69 | 3.61E-04 |
| CC | GO:0031253 | cell projection membrane | 92 | 4.34E-04 |
| CC | GO:0031252 | cell leading edge | 108 | 5.26E-04 |
| MF | GO:0019787 | ubiquitin-like protein transferase activity | 106 | 2.20E-05 |

GO: Gene Ontology; DEGs: differentially expressed genes; BP: biological process;
CC: cell component; MF: molecular function; AD-MSCs: adipose-derived mesenchymal stem cells.

### 3.3 Signaling pathway enrichment analysis

KEGG pathway analysis revealed that the 456 overlapped DEGs were significantly enriched in the cell proliferation related pathways, such as cell cycle, DNA replication, and oocyte meiosis (Fig. 2A). For 6 specifically upregulated DEGs in AD-MSCs, OLR1 gene was enriched in PPAR signaling pathway and phagosome, while INHBA gene was enriched in 3 KEGG pathways, including TGF-beta signaling pathway, signaling pathways regulating pluripotency of stem cells, and cytokine-cytokine receptor interaction. No significant pathway enrichment was found in the rest four genes (Fig. 2B). The top 10 most significant pathways of the specifically downregulated DEGs in AD-MSCs are shown in Fig. 2C. These downregulated genes were mainly involved in the metabolic pathways, pathways in cancer, and infection-related pathways (hepatitis C, hepatitis B, herpes simplex infection, and measles). For BM-MSCs specific DEGs in GSE36474, the top 10 terms of KEGG category are presented in Supplementary Fig. 1.

**Fig. 2.**
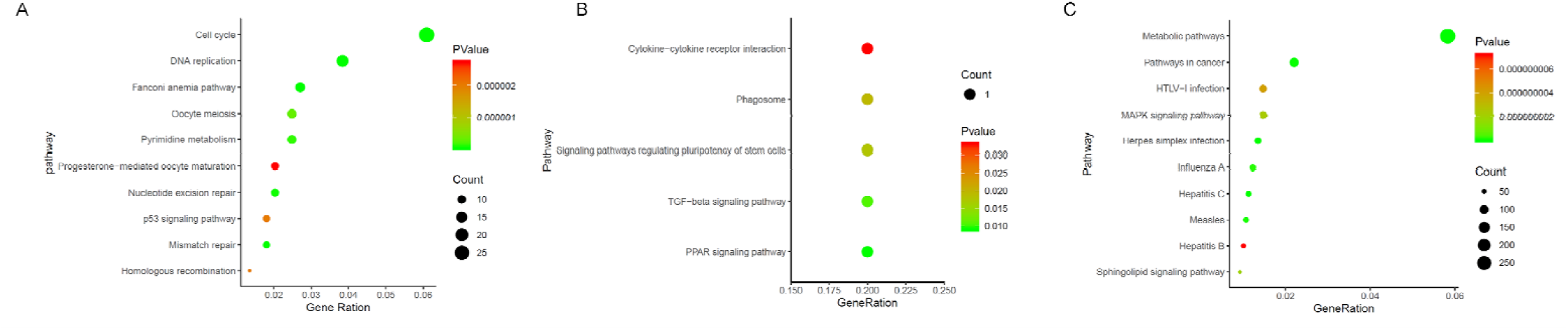
The Kyoto Encyclopedia of Genes and Genomes (KEGG) pathway analysis of differentially expressed genes (DEGs): (A) The top 10 significant KEGG pathways of common downregulated DEGs. (B) KEGG pathway enrichment analysis of specifically upregulated DEGs in adipose-derived mesenchymal stem cells (AD-MSCs). (C) The top 10 significant KEGG pathways of specifically downregulated DEGs in AD-MSCs.

### 3.4 PPI network analysis of the DEGs and the identification of hub genes

The PPI network of 456 overlapped DEGs was constructed. After removing the genes that were not connected in the network, a total of 389 nodes and 1936 edges were obtained (Fig. 3A). Seven overlapped genes (i.e., CDK1, BUB1B, CCNB1, CDC20, MAD2L1, NDC80, and CDCA8) were identified as hub genes by the three ranking methods (Supplementary Fig. 2). Furthermore, the PPI network of seven hub genes was constructed by STRING to further understand the interaction of these genes and the results showed that they are closely associated with each other (Supplementary Fig. 3). According to the degree of importance, we chose 3 significant modules from the PPI network complex for further analysis using Cytotype MCODE. Module 1 consisted of 26 nodes and 325 edges (Fig. 3B, Table 3), which were markedly enriched in cell cycle and oocyte meiosis. Module 2 consisted of 20 nodes and 165 edges (Fig. 3C, Table 3), which were mainly associated with cell cycle. Module 3 consisted of 31 nodes and 236 edges (Fig. 3D, Table 3), which were mostly enriched in DNA replication, cell cycle and fanconi anemia pathway. In addition, all 7 hub genes were clustered in Module 1 and most of the hub genes were associated with cell cycle pathway. These results suggested that the cell cycle pathway plays significant roles in MM. A heat map was employed to present the differences in the expression levels of the 7 hub genes in GSE133346 and GSE36474, respectively (Fig. 4). The results revealed that these genes were significantly downregulated in both AD-MSCs and BM-MSCs from MM patients.

**Table 3.**
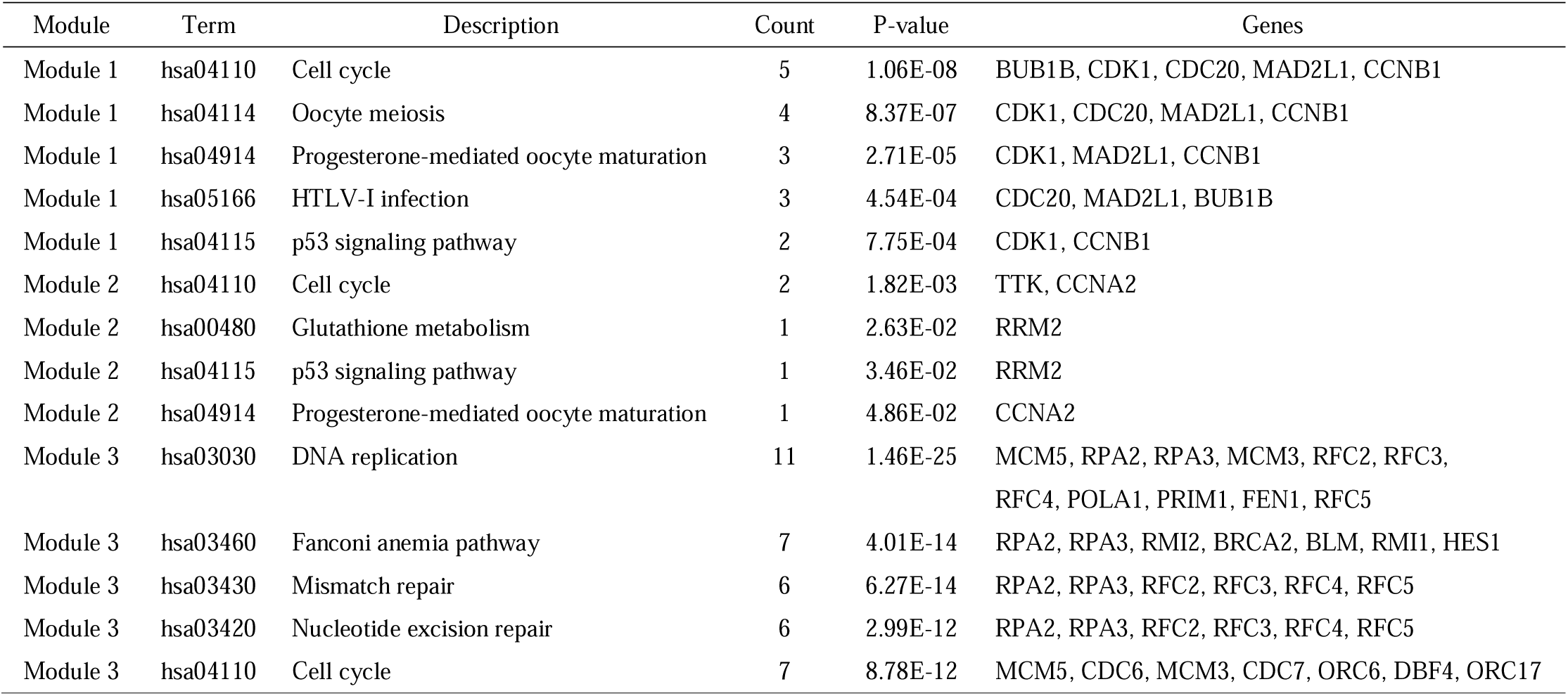
Kyoto Encyclopedia of Genes and Genomes (KEGG) pathway enrichment analysis of the differentially expressed genes (DEGs) in top 3 modules.

| Module | Term | Description | Count | P-value | Genes |
| --- | --- | --- | --- | --- | --- |
| Module 1 | hsa04110 | Cell cycle | 5 | 1.06E-08 | BUB1B, CDK1, CDC20, MAD2L1, CCNB1 |
| Module 1 | hsa04114 | Oocyte meiosis | 4 | 8.37E-07 | CDK1, CDC20, MAD2L1, CCNB1 |
| Module 1 | hsa04914 | Progesterone-mediated oocyte maturation | 3 | 2.71E-05 | CDK1, MAD2L1, CCNB1 |
| Module 1 | hsa05166 | HTLV-I infection | 3 | 4.54E-04 | CDC20, MAD2L1, BUB1B |
| Module 1 | hsa04115 | p53 signaling pathway | 2 | 7.75E-04 | CDK1, CCNB1 |
| Module 2 | hsa04110 | Cell cycle | 2 | 1.82E-03 | TTK, CCNA2 |
| Module 2 | hsa00480 | Glutathione metabolism | 1 | 2.63E-02 | RRM2 |
| Module 2 | hsa04115 | p53 signaling pathway | 1 | 3.46E-02 | RRM2 |
| Module 2 | hsa04914 | Progesterone-mediated oocyte maturation | 1 | 4.86E-02 | CCNA2 |
| Module 3 | hsa03030 | DNA replication | 11 | 1.46E-25 | MCM5, RPA2, RPA3, MCM3, RFC2, RFC3, RFC4, POLA1, PRIM1, FEN1, RFC5 |
| Module 3 | hsa03460 | Fanconi anemia pathway | 7 | 4.01E-14 | RPA2, RPA3, RMI2, BRCA2, BLM, RMI1, HES1 |
| Module 3 | hsa03430 | Mismatch repair | 6 | 6.27E-14 | RPA2, RPA3, RFC2, RFC3, RFC4, RFC5 |
| Module 3 | hsa03420 | Nucleotide excision repair | 6 | 2.99E-12 | RPA2, RPA3, RFC2, RFC3, RFC4, RFC5 |
| Module 3 | hsa04110 | Cell cycle | 7 | 8.78E-12 | MCM5, CDC6, MCM3, CDC7, ORC6, DBF4, ORC17 |

**Fig. 3.**
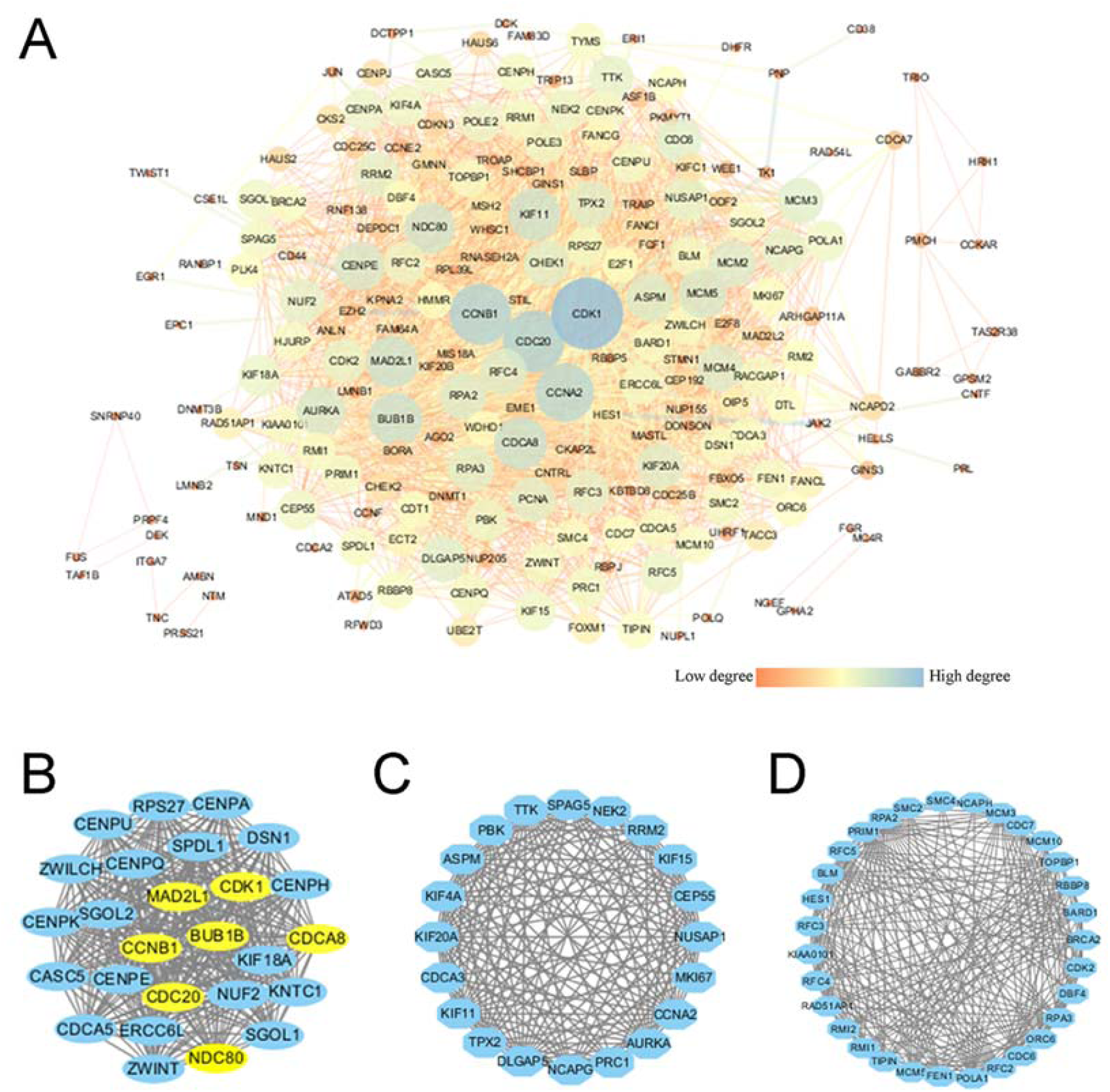
The protein-protein interaction (PPI) network and modules identified: (A) PPI network analysis of overlapped differentially expressed genes (DEGs). Nodes with higher degree are displayed in larger size and dark blue color and nodes with lower degree are shown in smaller size and bright orange color. The edge size is consistent with the coexpression intensity. Modules 1 (B), Modules 2 (C) and Modules 3 (D) were identified in PPI. The blue circle indicates the downregulated DEGs, the top 7 hub genes are highlighted with yellow circles.

**Fig. 4.**
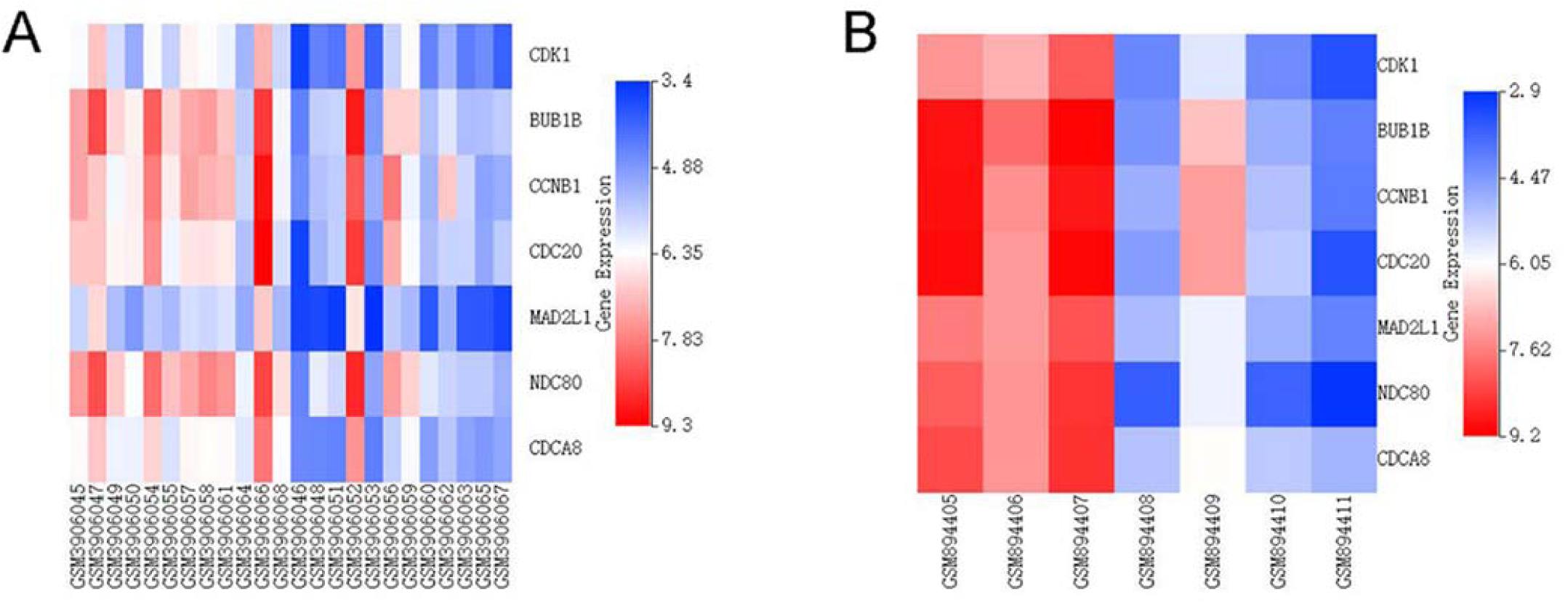
Hub gene expression heat maps among the 2 databases: (A) GSE133346, (B) GSE36474. Red color represents high expression level, and blue color represents low expression level.

## 4 Discussion

MSCs play important roles in MM pathogenesis. However the exact molecular mechanism of MSCs in MM development and progression remains unclear. Especially for AD-MSCs, no previous study of the bioinformatics analysis was conducted to explore the key genes and pathways of AD-MSCs in MM to the best of our knowledge. Therefore, the expression of genes in two microarray datasets based on MSCs (AD-MSCs and BM-MSCs) of MM patients and their normal counterparts were analyzed in current study. A total of 456 common downregulated DEGs in AD-MSCs and BM-MSCs were identified. In addition, 6 specifically upregulated and 5143 specifically downregulated DEGs in GSE133346 were further identified as specific DEGs of AD-MSCs, while 139 specifically upregulated and 537 specifically downregulated DEGs in GSE36474 were further identified as specific DEGs of BM-MSCs. We also performed GO and KEGG pathway analysis to find crucial pathways. Furthermore, the PPI network of 456 common downregulated DEGs was constructed and 7 overlapped genes were identified as hub genes by the three ranking methods, which increased the reliability of the results compared with the method of screening hub genes by degree. The present study may help us form a comprehensive insight into the mechanisms of MM and identify potential biomarkers for prognosis and therapy of MM.

In this study, seven hub genes were identified, most of which were reported to be related to the development and progression of MM [8, 12, 13]. However, it is noteworthy that few studies directly addressed the potential effects of these genes in MSCs from MM patients. Among these hub genes, the BUB1B gene and its encoded protein (BUBR1) were reported to serve as a checkpoint for proper chromosome segregation and to prevent separation of the duplicated chromosomes in normal cells [14-16]. Previous investigations indicated that increased BUB1B expression in MM cells demonstrated drug-resistance feature, promoted MM cell proliferation and led to poor survival in MM patients [14]. However, it was interesting that, in this study, BUB1B was downregulated in both AD- and BM-MSCs from MM patients. Previous study reported that targeted inhibition of BUBR1 expression in MSCs significantly reduced the potential to differentiate and led to cellular senescence [17]. On one hand, for AD-MSCs, myeloma cells can corrupt senescent AD-MSCs and impair their anti-tumor activity [18]. On the other hand, for BM-MSCs, senescent cells were also reported to exacerbate the progression of MM [11, 19]. Therefore, cellular senescence of both AD-MSCs and BM-MSCs may increase the proliferation of MM cell and led to the deterioration of the disease. These findings might reveal a novel function of BUB1B in MSCs and targeting BUB1B might be a potential therapeutic strategy for MM patients.

Moreover, as a cell proliferation-associated gene, CDK1 was reported to be an important driver for MM disease progression [20, 21]. Besides, previous studies also showed that the downregulation of CDK1 was related to osteogenic differentiation disorder and senescence of MSCs [22-24]. In addition, it was interesting that expression knockdown of CDK1 resulted in significantly increased adipogenic differentiation of AD-MSCs [25]. Therefore, abnormal expression of CDK1 in MSCs may also result in dysfunction of MSCs and lead to progression of MM although further functional studies are still needed.

As for MAD2L1, CCNB1, CDCA8, CDC20 and NDC80, these hub genes also played important roles in regulating cell proliferation, apoptosis and chromosome stability according to pervious reports [26, 27]. The irregular expression of these genes in cells often accompanied with cell proliferation disorders and chromosome instability [28-30]. As it was reported that genome instability and low proliferative capacity were key causes of MSCs senescence [24, 31, 32], downregulation of these identified hub genes in MSCs may also eventually lead to the progression of MM.

Importantly, according to KEGG pathway analysis, we found that common downregulated DEGs, the genes in top three modules, and the hub genes were all primarily enriched in the cell cycle that was documented previously as a factor to start the process of MM and was closely related to patients’ survival [13]. These data were in accordance with previous studies, which indicated that aberrant cell cycle pathway of MSCs marked with S phase arrest and reduced proliferation of MSCs played crucial roles in development of MM [11] and other cancers [33]. In addition, previous study also found that the function of cell cycle of MSCs was impaired in MM patients, leading the MSCs to enter into an early cellular senescence process [9] and to participate in progression and relapse of MM [11, 18]. Therefore, impaired cell cycle pathway of MSCs contributed directly or indirectly to the pathophysiological progress of MM. Further unravelling of the molecular mechanism about this pathway and relevant genes may be necessary for future efforts.

In addition, the present analyses identified 6 DEGs that were specifically upregulated in AD-MSCs and may serve as promising diagnostic and therapeutic targets in MM. However, it is of note that the effects and potential mechanisms of these genes in MSCs were rarely investigated so far. A small number of studies demonstrated the important roles of INHBA and OLR1 in MSCs. INHBA was a key regulator in the maintenance of pluripotency of stem cells [34], including AD-MSCs [35]. Previous study showed that INHBA overexpression was related to high chondrogenic and osteogenic potential of MSCs [34], which was further suggested to suppress proliferation of MM plasma cells [36]. Therefore, upregulation of INHBA in AD-MSCs may also enhance osteogenic differentiation through osteogenic pathway (TGF-beta signaling pathway) and pluripotency-regulating pathway of stem cells, which might exert anti-tumor effects in MM and may be a promising therapeutic target in MM. As for OLR1 gene, it is a novel PPAR-γ target gene playing a critical role in the regulation of adipocyte lipid metabolism [37]. High level of OLR1 expression was linked to obesity and dyslipidemia [37-39]. PPAR-γ was well established as a prime inducer of adipogenesis, which was demonstrated to regulate the differentiation of MSCs into adipocytes [40]. Based on these previous findings, overexpressed OLR1 in AD-MSCs might also result in increased adipogenic differentiation by PPAR signaling pathway, which would eventually enhance MM cell growth [41]. As for the rest 4 identified DEGs (i.e., GPR116, BEX1, AMIGO2 and CNN1), it is interesting that currently no studies were found to investigate their roles in AD-MSCs so far as we know. These genes might hence be novel potential diagnostic and therapeutic biomarkers in MM although further functional research is still required.

Furthermore, this study also identified 5143 specifically downregulated DEGs in AD-MSCs. Among these genes, IL-1R2 gene (IL1R2) and ZNF45 were demonstrated to be expressed uniquely in AD-MSCs but not in BM-MSCs [42]. Anti-inflammatory cytokine IL-1R2 is a specific and natural inhibitor of IL-1 and was reported to regulate cell metabolism and to respond to immune inflammation induced by many cytokines [43]. Previous studies showed that IL-1 inhibited chondrogenesis of adult AD-MSCs [44]. Therefore, downregulated IL1R2 in AD-MSCs may weaken its inhibitory effect on IL-1 and result in chondrogenic disorder of AD-MSCs and development of MM. As for ZNF45, this gene is a zinc finger gene, which was involved in transcription control in AD-MSCs [42]. Previous studies demonstrated that many zinc finger genes/proteins functioned as key transcriptional regulators in osteoblast differentiation of MSCs [45-47]. So, changes in the expression of ZNF45 in AD-MSCs may also lead to dysfunction of osteogenic differentiation of AD-MSCs. These two genes may be important markers of AD-MSCs in MM patients and promising therapeutic targets for MM. Further research on the underlying molecular mechanisms of these two genes in AD-MSCs may supply new insights for the treatment of MM.

In conclusion, through a series of comprehensive analysis of bioinformatics, several crucial genes and certain associated pathways were successfully screened. Our study suggested that the cell cycle pathway was closely related to the development of MM. Seven hub genes may be the potential diagnostic and therapeutic biomarkers both in AD-MSCs and BM-MSCs from MM patients. In addition, IL1R2 and ZNF45 might act as important markers of AD-MSCs in MM. These findings would provide a comprehensive insight into the pathogenesis of MM and hold promise for finding potential drug targets and diagnostic biomarkers.

## Author Contributions

Study conceive and design (ZRL, WKC, GHH), acquisition of subjects and data (ZRL, WKC, GHH), analysis and interpretation of data (ZRL, WKC, QY, MLS, HZZ, QH, GXZ, KYC, GHH), and preparation of manuscript (ZRL, WKC, QY, MLS, HZZ, QH, GXZ, KYC, GHH).

## Acknowledgements

Not applicable.

## Founding

This work was supported by grants from the National Science Foundation of China (NO. 81460560 and 81960664) and the Applied Basic Research Program of Yunnan Province of China (NO. 2017FB134).

