## Supplementary Table 1 for "Identification of potential biomarkers or therapeutic targets of mesenchymal stem cells in multiple myeloma by bioinformatics analysis"

Supplementary Material Table 1. GO enrichment analysis of the specifically up- and downregulated DEGs in BM-MSCs.

| Category | Term | Description | Count | P value |
|---|---|---|---|---|
| Upregulated | | | | |
| BP | GO:0060337 | type I interferon signaling pathway | 10 | 1.78E-10 |
| BP | GO:0071357 | cellular response to type I interferon | 10 | 1.78E-10 |
| BP | GO:0034340 | response to type I interferon | 10 | 2.74E-10 |
| BP | GO:0009615 | response to virus | 14 | 8.82E-09 |
| BP | GO:0051607 | defense response to virus | 12 | 1.49E-08 |
| BP | GO:0098542 | defense response to other organism | 12 | 3.31E-05 |
| BP | GO:0050688 | regulation of defense response to virus | 5 | 7.87E-05 |
| BP | GO:0022407 | regulation of cell-cell adhesion | 10 | 1.21E-04 |
| BP | GO:0043900 | regulation of multi-organism process | 10 | 1.79E-04 |
| BP | GO:1901342 | regulation of vasculature development | 10 | 2.48E-04 |
| MF | GO:0005539 | glycosaminoglycan binding | 7 | 2.51E-04 |
| MF | GO:0005125 | cytokine activity | 6 | 4.10E-04 |
| Downregulated | | | | |
| BP | GO:0071103 | DNA conformation change | 31 | 7.71E-14 |
| BP | GO:0007059 | chromosome segregation | 29 | 1.90E-12 |
| BP | GO:0032392 | DNA geometric change | 15 | 4.30E-10 |
| BP | GO:0006260 | DNA replication | 25 | 4.81E-10 |
| BP | GO:0048285 | organelle fission | 31 | 6.37E-10 |
| BP | GO:0000280 | nuclear division | 29 | 1.05E-09 |
| BP | GO:0000226 | microtubule cytoskeleton organization | 33 | 1.74E-09 |
| BP | GO:0006310 | DNA recombination | 23 | 3.55E-09 |
| BP | GO:0032508 | DNA duplex unwinding | 13 | 8.15E-09 |
| BP | GO:0006336 | DNA replication-independent nucleosome assembly | 11 | 1.40E-08 |
| CC | GO:0000793 | condensed chromosome | 29 | 2.96E-16 |
| CC | GO:0098687 | chromosomal region | 35 | 1.04E-14 |
| CC | GO:0000776 | kinetochore | 21 | 6.03E-13 |
| CC | GO:0000775 | chromosome, centromeric region | 24 | 1.55E-12 |
| CC | GO:0000779 | condensed chromosome, centromeric region | 18 | 1.94E-11 |
| CC | GO:0000777 | condensed chromosome kinetochore | 16 | 3.65E-10 |
| CC | GO:0005819 | spindle | 24 | 1.58E-07 |
| CC | GO:0000940 | condensed chromosome outer kinetochore | 5 | 3.04E-06 |
| CC | GO:0000794 | condensed nuclear chromosome | 10 | 4.25E-06 |
| CC | GO:0044450 | microtubule organizing center part | 13 | 7.64E-05 |
| MF | GO:0008094 | DNA-dependent ATPase activity | 12 | 8.13E-08 |
| MF | GO:0140097 | catalytic activity, acting on DNA | 15 | 2.31E-06 |
| MF | GO:0003678 | DNA helicase activity | 8 | 4.32E-06 |
| MF | GO:0000217 | DNA secondary structure binding | 6 | 1.03E-05 |
| MF | GO:0008017 | microtubule binding | 14 | 3.02E-05 |
| MF | GO:0004003 | ATP-dependent DNA helicase activity | 6 | 6.42E-05 |
| MF | GO:0003684 | damaged DNA binding | 8 | 6.52E-05 |

| | | | | |
|---|---|---|---|---|
| MF | GO:0015631 | tubulin binding | 16 | 2.51E-04 |
| MF | GO:0000400 | four-way junction DNA binding | 4 | 4.02E-04 |
| MF | GO:0004518 | nuclease activity | 12 | 6.07E-04 |

GO: Gene Ontology; DEGs: differentially expressed genes; BM-MSCs: bone marrow-derived mesenchymal stem cells.
