## Supplemental Figures for "Identification of potential biomarkers or therapeutic targets of mesenchymal stem cells in multiple myeloma by bioinformatics analysis"

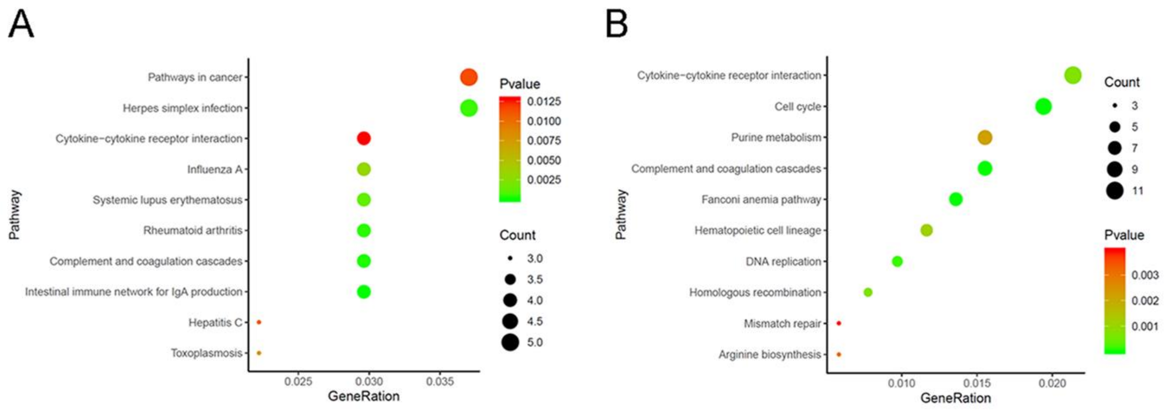


**Supplementary Material Fig. 1.** The top 10 KEGG pathway analysis of the specifically up- and downregulated DEGs in bone marrow-derived mesenchymal stem cells (BM-MSCs): (A) specifically upregulated DEGs, (B) specifically downregulated DEGs.

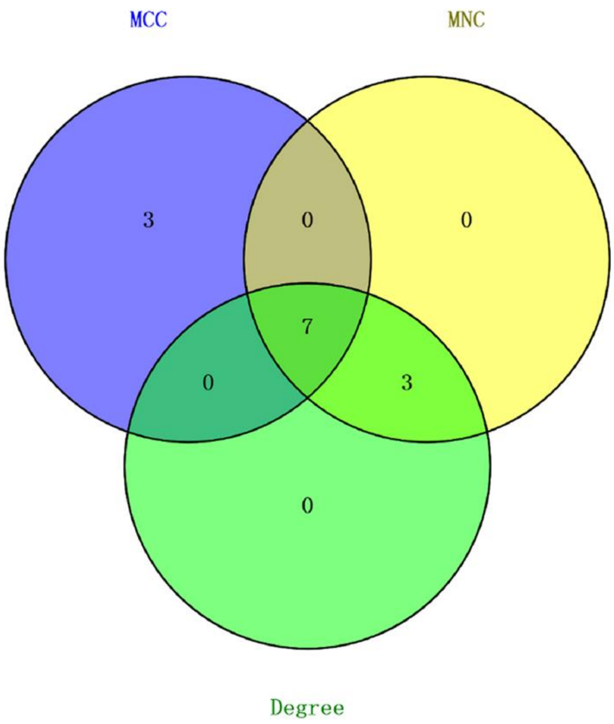


**Supplementary Material Fig. 2.** The Venn diagram of mutual key genes based on three methods. MCC: maximal clique centrality; MNC: maximum neighborhood component; Degree: the connectivity of the protein.

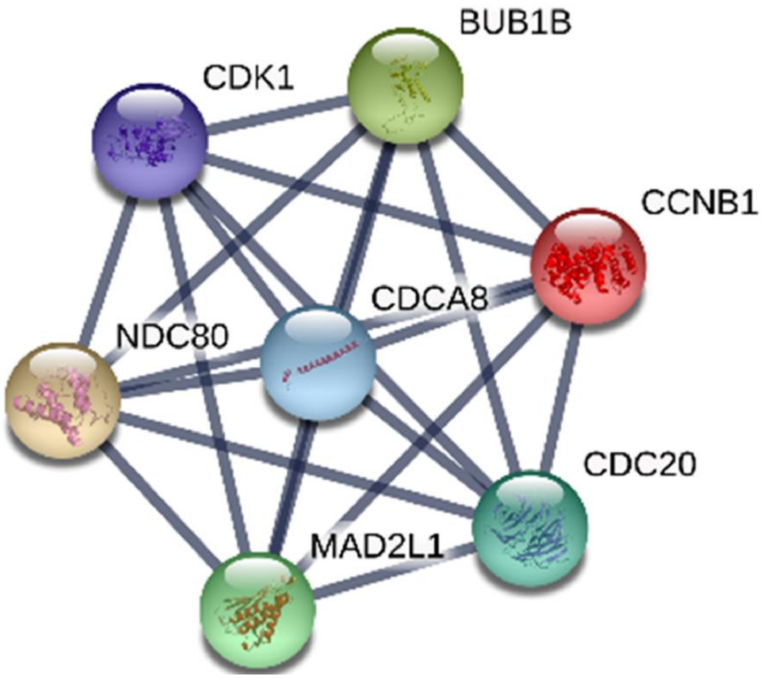


**Supplementary Material Fig. 3.** The protein-protein interaction (PPI) network analysis of 7 hub genes.
